# The influence of stimulus and behavioral histories on predictive control of smooth pursuit eye movements

**DOI:** 10.1101/2021.06.05.447182

**Authors:** Takeshi Miyamoto, Yutaka Hirata, Akira Katoh, Kenichiro Miura, Seiji Ono

**Affiliations:** Faculty of Health and Sport Sciences, University of Tsukuba, Ibaraki, Japan; Department of Robotic Science and Technology, Chubu University College of Engineering, Kasugai, Japan; Department of Physiology, Tokai University School of Medicine, Kanagawa, Japan; Department of Pathology of Mental Diseases, National Institute of Mental Health, National Center of Neurology and Psychiatry, Tokyo, Japan; Graduate School of Medicine, Kyoto University, Kyoto, Japan

## Abstract

The pursuit system has the ability to perform predictive control of eye movements. Even when the target motion is unpredictable due to velocity or direction changes, preceding changes in eye velocity are generated based on weighted averaging of past stimulus timing. However, it is still uncertain whether behavioral history influences the control of predictive pursuit. Thus, we attempted to clarify the influences of stimulus and behavioral histories on predictive pursuit to randomized target velocity. We used alternating-ramp stimuli, where the rightward velocity was fixed while the leftward velocity was either fixed (predictable) or randomized (unpredictable). Predictive eye deceleration was observed regardless of whether the target velocity was predictable or not. In particular, the predictable condition showed that the predictive pursuit responses corresponded to future target velocity. The linear mixed-effects model showed that both stimulus and behavioral histories of the previous two or three trials had influences on the predictive pursuit responses to the unpredictable target velocity. Our results suggest that the predictive pursuit system allows to track randomized target motion using the information from previous several trials, and the information of sensory input (stimulus) and motor output (behavior) in the past time sequences have partially different influences on predictive pursuit.

## Introduction

Smooth pursuit eye movements allow us to continuously hold a moving object on or near the fovea, resulting in high visual acuity for the target. In general, the human oculomotor system requires approximately 100 ms from the onset of target motion to the onset of eye movement (Carl and Gellman 1987; Leigh and Zee 2015), and this delay can lead to a loss of visual information because visual acuity declines as the target moves from the fovea to the retinal periphery (Jacobs 1979). To overcome the inherent delay, the oculomotor system has the ability to perform predictive feedforward control of eye movements (Fiehler et al. 2019; Kowler et al. 2019). When the timing of onset/offset or directional changes of the target motion is predictable, preceding eye velocity is generated according to future visual motion (Barnes 2008; Fukushima et al. 2013). Such predictive pursuit responses are thought to be pre-programmed (Barnes et al. 2005; Barnes and Schmid 2002), and its timing and scale can be built up with repeated trials (Barnes and Schmid 2002; Chakraborti et al. 2002; Collins and Barnes 2005).

Predictive pursuit is known to occur even when the timing of onset/offset or directional change of visual motion is randomized, and the timing of onset of the predictive pursuit is influenced by the history of previous trials (Heinen et al. 2005; de Hemptinne et al. 2013). Several studies have demonstrated that the effect of stimulus history on the onset of predictive pursuit does not depend solely on the most recent trial, but is determined by sequence of several previous trials, like the branches of a tree (Badler and Heinen 2006; de Hemptinne et al. 2010). Moreover, other studies have demonstrated that the timing of onset of predictive pursuit to visual motion with the randomized timing of directional changes is quantitatively associated with the ramp duration of previous trials, and the influence of previous trials on the predictive pursuit declines progressively as the previous trial retracts into the past (Barnes and Collins 2011, 2015; Collins and Barnes 2009). Taken together, the stimulus information of previous trials appears to be continuously averaged and used to determine the timing of onset of predictive pursuit to visual motion with the randomized timing.

Eye velocity is an important component of predictive smooth pursuit, as well as the timing of onset. Although predictive eye velocity is regulated according to predictable velocity of future target motion (Boman and Hotson 1992; Heinen et al. 2005; Jarrett and Barnes 2002), it is still uncertain whether the strategy of weighted averaging of the past stimulus information is applied to predictive pursuit to target motion with the randomized velocity as it does for the randomized timing. For predictive pursuit of randomly timed stimuli, a model of ocular pursuit described by previous studies has proposed that a continual estimation of the timing of onset of future target motion is constructed by weighted averaging of stimulus history and retained in a form of working memory to initiate predictive pursuit (Barnes and Collins 2011; Fukushima et al. 2013). This model is supported by fMRI studies that show an activity of the dorsolateral prefrontal cortex (DLPFC), which is associated with working memory (Goldman-Rakic 1991), during smooth pursuit to predictable target motion (Burke and Barnes 2008a; Schmid et al. 2001). Considering the behavioral evidence that the velocity of predictive pursuit is built up with repeated trials (Barnes and Schmid 2002; Chakraborti et al. 2002; Collins and Barnes 2005), the information of target velocity may be retained in working memory as well. However, unlike the timing of onset, the velocity of predictive pursuit can also be modulated by open-loop gain. The frontal eye field (FEF) is known to play a role in the control of open-loop gain (Tanaka and Lisberger 2001). A recent study has demonstrated that the gain control of FEF is modulated by the combination of behavior on previous trials and current visual information during the initial phase of smooth pursuit (Darlington et al. 2018). It has been demonstrated that the smooth pursuit gain is updated for each trial depending on whether observers have performed smooth pursuit in previous trials or not (Tabata et al. 2008). Although the above studies are not about predictive pursuit, it is suggested that behavioral history influences the eye velocity of predictive pursuit. Therefore, we hypothesized that the stimulus (i.e., velocity of target motion) and behavioral (i.e., pursuit responses) histories affect the eye velocity of predictive pursuit, and this study was designed to clarify, in a quantitative sense, the influence of both histories on the predictive pursuit to visual motion with randomized velocity.

## Materials and Methods

### Observers

The observers were 12 adults (3 women, 9 men, mean age: 23.6 [SD: 1.3] years old) and they reported having normal or corrected to normal vision and no known visuomotor deficits. The observers were diagnosed neither with stereoscopic problem nor strabismus. All the observers gave written informed consent in accordance with the Declaration of Helsinki. All the protocols were approved by the Research Ethics Committee at the Faculty of Health and Sport Sciences, University of Tsukuba.

### Apparatus

The observers sat 57 cm in front of a CRT monitor (22-inch, RDF223G, Mitsubishi, refresh rate: 60 Hz, spatial resolution: 800 × 600 pixels, background luminance: 60 cd/m^2^) with head stabilized by a chin rest and a forehead restraint. Eye movements from the right eye were detected using a video-based eye tracking system (Matsuda et al. 2017; Miyamoto et al. 2020; Ono et al. 2019). The eye position signals detected by the system were digitized at 1 kHz with 16-bit precision using CED-Micro 1401 hardware (Cambridge Electronic Designs, Cambridge, England). Prior to the task, the eye position signals were calibrated by requiring the observers to fixate a target spot (diameter of 0.3 deg) at known horizontal and vertical eccentricities in binocular viewing condition. The target consisted of a white Gaussian dot (SD: 0.15 deg) on a uniform black background. All the visual stimuli were generated by Psychophysics Toolbox extensions on MATLAB (Mathworks, MA, US).

### Procedure

The target moved horizontally with constant velocity over a distance of 24 deg in an irregular triangular waveform (alternating-ramp). The rightward target velocity was always 16 deg/s, while the leftward velocity was one of seven (4, 8, 12, 16, 20, 24, 28 deg/s). The position at which the target would reverse was always indicated to the observers by arrows. The observers were instructed to track the target as accurately as possible. The task consisted of predictable and unpredictable conditions, the order of which was randomized among the observers. In the predictable condition (Fig. 1 blue lines), one of the seven speeds was repeated 30 times within a block, the order of the seven velocities was randomized, and there was a one-minute interval between each block. In the unpredictable condition (Fig. 1 red lines), the seven velocities were presented randomly within one block. Each block for the unpredictable condition consisted of 49 trials (7 velocities × 7 trials), and 5 blocks were conducted with one-minute intervals. Prior to the task, the observers practiced by performing 21 trials following the presentation method of each condition.

**Fig. 1.**
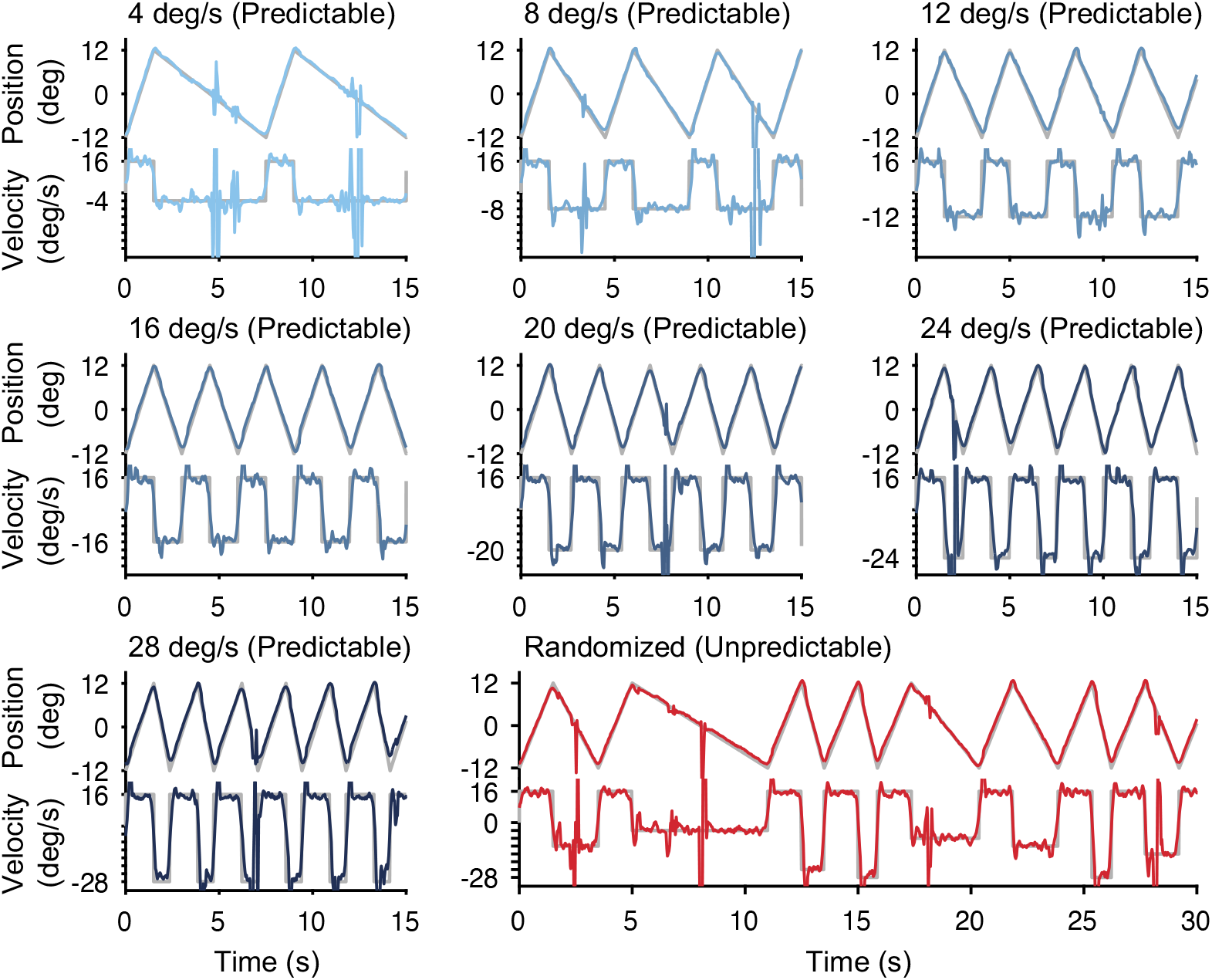
Triangular smooth pursuit of a small-diameter target moving horizontally. The target moved to the right at 16 deg/s and then reversed to the left at one of seven velocities. The upper part of each panel represents the target and eye position, and the lower part represents the target and eye velocity. The gray lines represent the target position and velocity. The blue lines represent the eye position and velocity in the predictable condition in which a single target velocity to the left was repeated within a block. The red lines represent the eye position and velocity in the unpredictable condition in which the target velocity to the left was randomized. Upward deflections show rightward eye motion.

### Eye movement recordings and analysis

Eye velocity and acceleration were generated by digitally differentiating the position arrays using the central difference algorithm in Matlab (Mathworks, MA, US). Velocity and acceleration data were filtered using an 80-point finite impulse response (FIR) digital filter with a 30 Hz passband. Saccades were identified according to a criterion of acceleration of 1000 deg/s and linear interpolation was applied to fill the gaps left by the removed saccades. Because oscillatory fluctuations remained in the eye velocity traces after the above filtering, a moving average was applied over a 40 ms window (Fig 2).

**Fig. 2.**
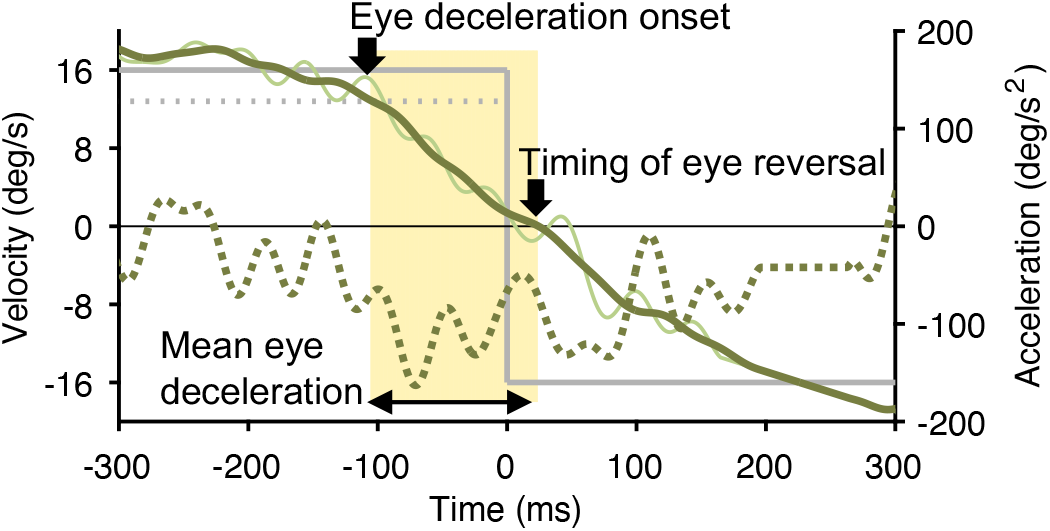
A sample trial near the timing of target reversal. The y-axis on the left represents the velocity, which corresponds to the eye velocity after filtering (30 Hz, light green solid line), the eye velocity with moving average applied (40-ms window, dark green solid line), the target velocity (gray solid line), and the threshold to detect the onset of predictive pursuit (gray dotted line). The y-axis on the right corresponds to the eye acceleration (dark green dotted line). The yellow shade represents the time that the mean eye deceleration was calculated. Time of 0 ms represents the timing of target reversal. Upward deflections show rightward eye motion.

The predictive pursuit for each trial was evaluated when the target reversed from the right to left. First, the eye deceleration onset was defined as the timing when the eye velocity fell below a threshold calculated as 80% of the average of the eye velocities of all the trials in the steady-state phase (700-500 ms before the target reversal). Second, the timing of eye reversal was defined as the timing when the eye velocity fell below 0 deg/s. Third, the mean eye deceleration was calculated as the average of eye acceleration between the eye deceleration onset and the timing of eye reversal. The above evaluation methods for predictive pursuit are summarized in Fig 2.

Trials in which the eye deceleration onset was earlier than −300 ms relative to the timing of target reversal were excluded. Most of the excluded trials involved a preceding saccade to the position where the target reverses. Based on this criteria, 139 (5.5%) out of a total of 2520 trials for the predictable condition and 179 (6.1%) out of a total of 2940 trials for the unpredictable condition were excluded. In addition, the first five trials of each block were removed and then the mean value of each variable was calculated according to the target velocity in both conditions.

## Results

### Predictable condition

Figure 3 shows averaged traces of pursuit eye velocity for each target velocity from a representative observer. The eye velocity started to decay prior to the target reversal, indicating that the observer performed predictive pursuit for the future target motion. One-way analyses of variance (ANOVA) with repeated measures using the target velocity in the left direction as the independent variable showed that all of the eye deceleration onset, the timing of eye reversal, and the mean eye deceleration varied according to the target velocity (the eye deceleration onset: *F*_6,66_ = 9.04, *p* = 3.29 × 10^−7^, partial *η*^2^ = 0.45, Fig. 4A; the timing of eye reversal: *F*_6,66_ = 50.59, *p* = 8.34 × 10^−23^, partial *η*^2^ = 0.82, Fig. 4B; the mean eye deceleration: *F*_6,66_ = 3.47, *p* = 4.8 × 10^−3^, partial *η*^2^ = 0.24, Fig. 4C). Moreover, within-subjects correlation coefficient, which is a method to focus on the changes of variable within each observer (Bland and Altman 1995), showed that the timing of eye reversal was associated closely with the target velocity (*r* = −0.90, *p* = 1.38 × 10^−27^) compared to the eye deceleration onset (*r* = −0.67, *p* = 1.33 × 10^−10^) and the mean eye deceleration (*r* = −0.46, *p* = 4.49 × 10^−5^). It appears that the predictive pursuit for the predictable target velocity is well reflected in the timing of eye reversal that is the result of control of both the eye deceleration onset and the mean eye deceleration. To corroborate this, we fitted a liner mixed-effects model of the timing of eye reversal with the eye deceleration onset and the mean eye deceleration as fixed effects, and individual intercept as the random effect [formula: the timing of eye deceleration ~ the eye deceleration onset + the mean eye deceleration + (1|observer)]. The result showed that both the eye deceleration onset and the mean eye deceleration had the significant fixed effects on the timing of eye deceleration (the eye deceleration onset: estimate ± SE = 0.690 ± 0.011, *t_2374_* = 63.90, *p* < 0.01; the mean eye deceleration: estimate ± SE = 0.595 ± 0.011, *t_2374_* = 55.10,*p* < 0.01), and the adjusted coefficient of determination (adjusted R^2^) was 0.68.

**Fig. 3.**
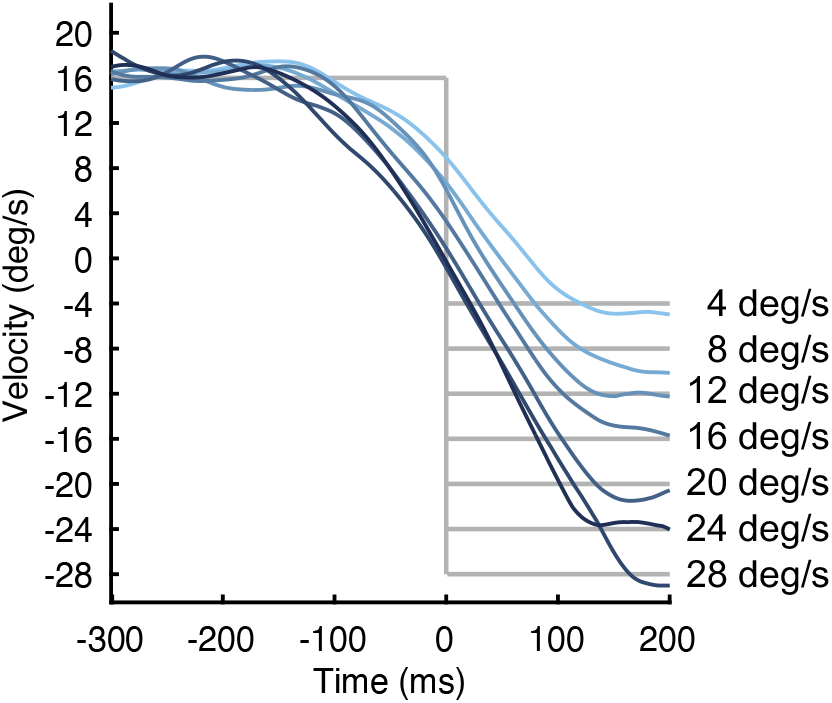
Averaged traces of pursuit eye velocity for each target velocity in the predictable condition from a representative observer. The colors of each eye velocity trace correspond to the seven target velocities (same as Fig. 1) and gray lines represent the target velocities. Time of 0 ms represents the timing of target reversal. Upward deflections show rightward eye motion.

**Fig. 4.**
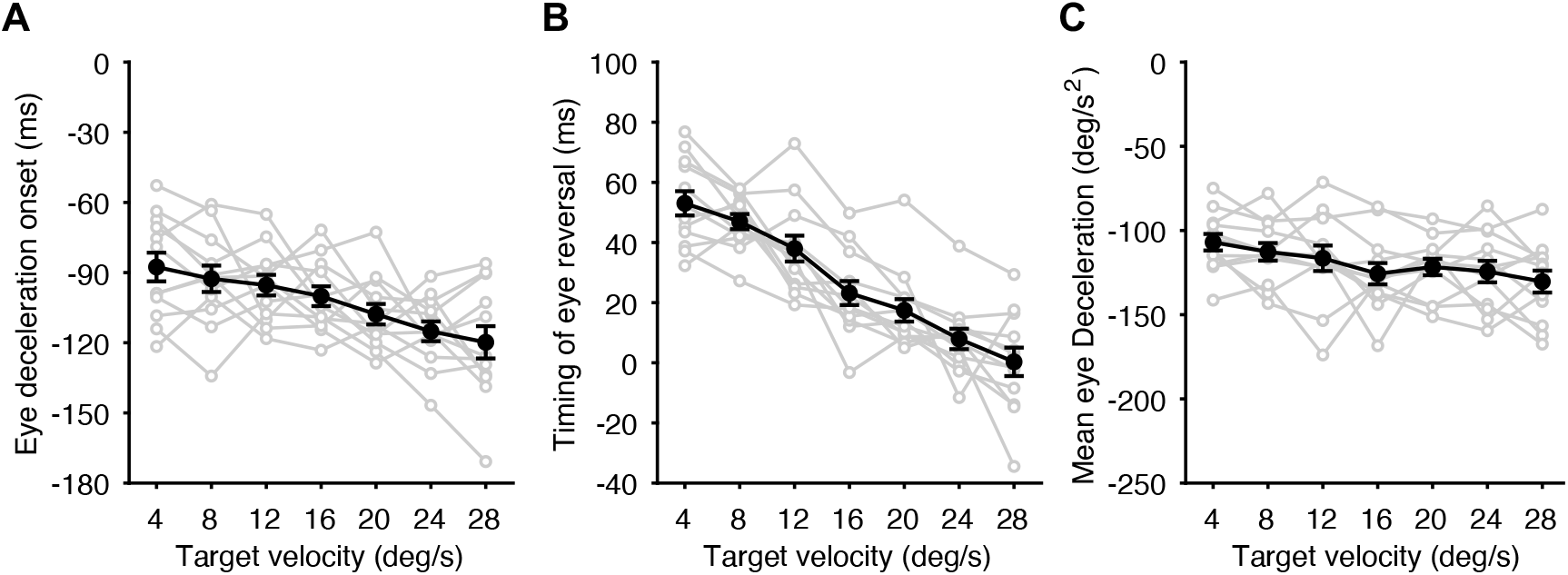
Predictive pursuit responses for the predictable condition. A: Eye deceleration onset relative to the target reversal as a function of target velocity. B: Timing of eye reversal relative to the target reversal as a function of target velocity. C: Mean eye deceleration as a function of target velocity. The gray circles and lines represent values of each observer and black circles and lines represent mean values of all the observers. Error bars indicate 1SE.

### Unpredictable condition

Predictive pursuit responses were observed even in the unpredictable condition, but they did not correspond to the future target velocity (the eye deceleration onset: *F*_6,66_ = 1.93, *p* = 0.09, partial *η*^2^ = 0.15, Fig. 5A; the timing of eye reversal: *F*_6,66_ = 1.64, *p* = 0.15, partial *η*^2^ = 0.13, Fig. 5B; the mean eye deceleration: *F*_6,66_ = 0.54, *p* = 0.78, partial *η*^2^ = 0.05, Fig. 5C). As with the predictable condition, the linear mixed-effects model showed that the timing of eye reversal was determined by two other indices (the eye deceleration onset: estimate ± SE = 0.627 ± 0.011, *t_2483_* = 55.98, *p* < 0.01; the mean eye deceleration: estimate ± SE = 0.474 ± 0.010, *t_2483_* = 46.89, *p* < 0.01. adjusted R^2^ = 0.62). Therefore, to examine the influence of stimulus and behavioral histories on the predictive pursuit to visual motion with randomized velocity, we used the timing of eye reversal as the representative index of predictive pursuit. A total of 15 linear mixed-effects models were set up with the timing of eye reversal in the current trial as the dependent variable and the target velocity (stimulus) and the timing of eye reversal (behavior) from the most recent trial (n-1) to before the 5th trial (n-5) as the independent variables, and individual intercept as the random effect. These models included those in which each history alone was incorporated into the model and those in which both were incorporated. Based on the Akaike’s information criterion (AIC), the model that incorporated the stimulus and behavioral histories up to before the 5th trial was adopted (Table 1 Model 15). In the model, the significant fixed effects were found for the stimulus history of the n-2 (estimate ± SE = −0.423 ± 0.085, *t_2447_* = −4.98, *p* = 6.8 × 10^−7^) and n-3 trials (estimate ± SE = −0.215 ± 0.085, *t_2447_* = −2.52, *p* = 0.01), and for the behavioral history of the n-1 (estimate ± SE = 0.086 ± 0.019, *t_2447_* = 4.52, *p* = 6.4 × 10^−6^) and n-2 trials (estimate ± SE = 0.101 ± 0.019, *t_2447_* = 5.41, *p* = 7.0 × 10^−8^).

**Fig. 5.**
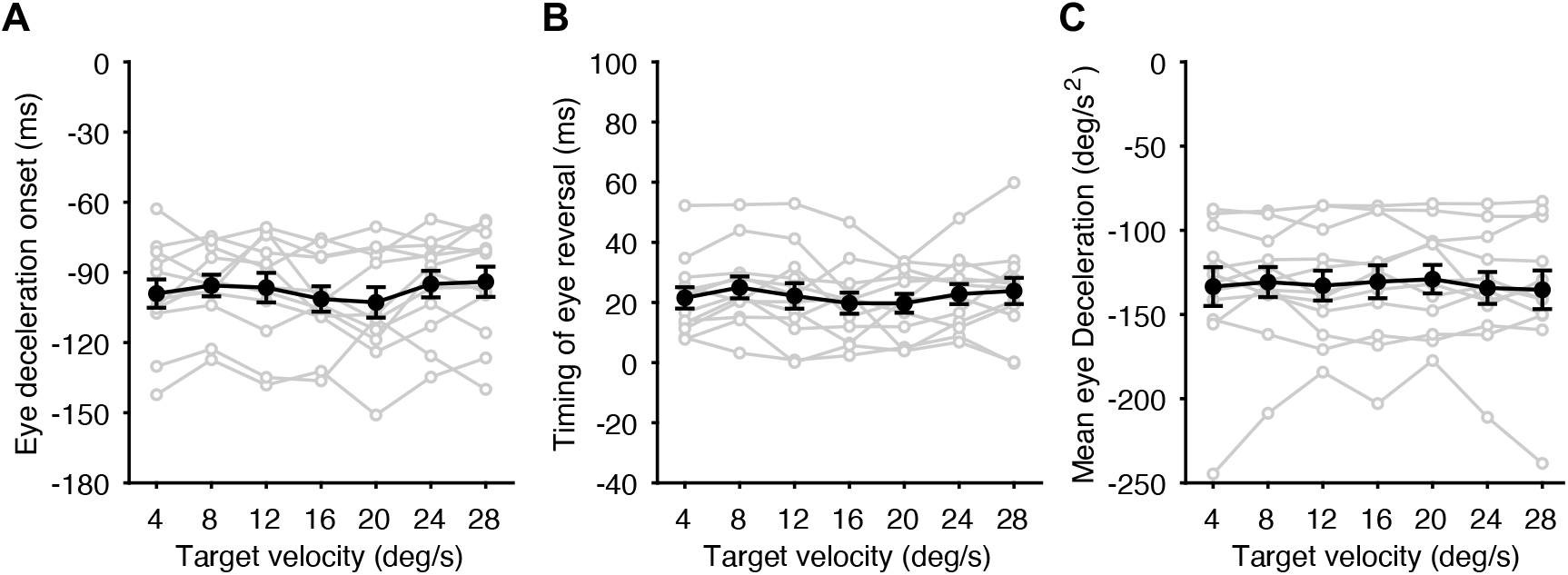
Predictive pursuit responses for the unpredictable condition. A: Eye deceleration onset relative to the target reversal as a function of target velocity. B: Timing of eye reversal relative to the target reversal as a function of target velocity. C: Mean eye deceleration as a function of target velocity. The gray circles and lines represent values of each observer and black circles and lines represent mean values of all the observers. Error bars indicate 1SE.

**Table 1.** Linear mixed-effects models with the timing of eye reversal in the current trial (n) for the unpredictable condition

|  | Model 1 |  | Model 2 |  | Model 3 |  | Model 4 |  | Model 5 |  |
| --- | --- | --- | --- | --- | --- | --- | --- | --- | --- | --- |
| Variable | Estimate | SE | Estimate | SE | Estimate | SE | Estimate | SE | Estimate | SE |
| (Intercept) | 24.633 | 3.488 | 31.357 | 3.841 | 35.745 | 4.189 | 36.330 | 4.502 | 37.687 | 4.766 |
| Stimulus (n-1) | -0.023 | 0.082 | -0.094 | 0.083 | -0.100 | 0.083 | -0.099 | 0.083 | -0.097 | 0.083 |
| Stimulus (n-2) |  |  | **-0.348** | **0.084** | **-0.398** | **0.086** | **-0.399** | **0.086** | **-0.395** | **0.086** |
| Stimulus (n-3) |  |  |  |  | **-0.219** | **0.084** | **-0.225** | **0.086** | **-0.227** | **0.086** |
| Stimulus (n-4) |  |  |  |  |  |  | -0.030 | 0.084 | -0.045 | 0.086 |
| Stimulus (n-5) |  |  |  |  |  |  |  |  | -0.073 | 0.084 |
| AIC | 24534.518 |  | 24519.286 |  | 24514.472 |  | 24516.346 |  | 24517.596 |  |

|  | Model 6 |  | Model 7 |  | Model 8 |  | Model 9 |  | Model 10 |  |
| --- | --- | --- | --- | --- | --- | --- | --- | --- | --- | --- |
| Variable | Estimate | SE | Estimate | SE | Estimate | SE | Estimate | SE | Estimate | SE |
| (Intercept) | 21.897 | 2.948 | 19.448 | 2.635 | 18.744 | 2.568 | 18.479 | 2.566 | 18.152 | 2.555 |
| Behavior (n-1) | **0.099** | **0.019** | **0.089** | **0.018** | **0.091** | **0.019** | **0.089** | **0.019** | **0.088** | **0.019** |
| Behavior (n-2) |  |  | **0.106** | **0.018** | **0.102** | **0.018** | **0.102** | **0.019** | **0.100** | **0.019** |
| Behavior (n-3) |  |  |  |  | 0.032 | 0.019 | 0.030 | 0.019 | 0.027 | 0.019 |
| Behavior (n-4) |  |  |  |  |  |  | 0.018 | 0.019 | 0.016 | 0.019 |
| Behavior (n-5) |  |  |  |  |  |  |  |  | 0.025 | 0.019 |
| AIC | 24420.771 |  | 24298.858 |  | 24230.987 |  | 24172.694 |  | 24119.207 |  |

|  | Model 11 |  | Model 12 |  | Model 13 |  | Model 14 |  | Model 15 |  |
| --- | --- | --- | --- | --- | --- | --- | --- | --- | --- | --- |
| Variable | Estimate | SE | Estimate | SE | Estimate | SE | Estimate | SE | Estimate | SE |
| (Intercept) | 22.197 | 3.218 | 27.009 | 3.325 | 30.637 | 3.685 | 30.015 | 4.063 | 30.154 | 4.373 |
| Stimulus (n-1) | -0.019 | 0.081 | -0.093 | 0.082 | -0.093 | 0.082 | -0.086 | 0.082 | -0.088 | 0.082 |
| Stimulus (n-2) |  |  | **-0.380** | **0.083** | **-0.433** | **0.085** | **-0.427** | **0.085** | **-0.423** | **0.085** |
| Stimulus (n-3) |  |  |  |  | **-0.214** | **0.083** | **-0.214** | **0.085** | **-0.215** | **0.085** |
| Stimulus (n-4) |  |  |  |  |  |  | 0.013 | 0.083 | 0.002 | 0.085 |
| Stimulus (n-5) |  |  |  |  |  |  |  |  | -0.017 | 0.083 |
| Behavior (n-1) | **0.099** | **0.019** | **0.090** | **0.018** | **0.089** | **0.019** | **0.087** | **0.019** | **0.086** | **0.019** |
| Behavior (n-2) |  |  | **0.106** | **0.018** | **0.104** | **0.018** | **0.103** | **0.019** | **0.101** | **0.019** |
| Behavior (n-3) |  |  |  |  | 0.032 | 0.018 | 0.030 | 0.019 | 0.027 | 0.019 |
| Behavior (n-4) |  |  |  |  |  |  | 0.015 | 0.019 | 0.014 | 0.019 |
| Behavior (n-5) |  |  |  |  |  |  |  |  | 0.022 | 0.018 |
| AIC | 24422.717 |  | 24281.751 |  | 24208.754 |  | 24153.043 |  | 24101.978 |  |

Dependent variable: the timing of eye reversal in the current trial.
Significant fixed effects are in bold ( $p < 0.05$ ).
Model 15 (including stimulus and behavioral histories from n-1 to n-5; the colored area) is adopted based on the AIC.

## Discussion

The purpose of this study was to clarify the influences of stimulus and behavioral histories on the predictive pursuit to target motion with randomized velocity using triangular target motion (alternating-ramp) stimuli. The target moved to the right at 16 deg/s and then reversed to the left at one of seven velocities. For the predictable condition in which a single target velocity to the left was repeated within a block, the predictive pursuit (eye deceleration) preceded the target reversal, as in previous studies (Barnes and Schmid 2002; Boman and Hotson 1992; Heinen et al. 2005; Kao and Morrow 1994). The eye deceleration onset and the mean eye deceleration varied dependent on the target velocity, and the timing of eye reversal reflected these predictive pursuit responses. Although predictive pursuit was observed even in the unpredictable condition in which the target velocity to the left was randomized, the responses did not correspond to the future target velocity. The predictive pursuit responses in the unpredictable condition appeared to converge to the intermediate values observed in the predictable condition. Such a centering strategy has been shown in previous studies using velocity randomization (Heinen et al. 2005) and timing randomization (Collins and Barnes 2009), and it appears to be effective in preventing large errors between eye and target motion in unpredictable situations while compensating for the delay in inherent visuomotor delays. To test the influence of stimulus and behavioral histories on predictive pursuit in the unpredictable condition, we compared different combinations of linear mixed-effects models. As a result, the model that incorporated the stimulus and behavioral histories up to last 5th trial was adopted, indicating both histories have fixed effects on the predictive pursuit in the current trial (n). Significant fixed effects were found for the stimulus history of the n-2 and n-3 trials (both estimates are negative values), and for the behavioral history of the n-1 and n-2 trials (both estimates are positive values). The signs of these fixed effects indicate that “the greater the target velocity in the previous trials” and “the greater the self’s predictive pursuit responses in the previous trials,” the greater the predictive pursuit response in the current trial, consistent with the results of previous studies that evaluated the history effect (Badler and Heinen 2006; Collins and Barnes 2009). Here, we discuss the underlying mechanisms involved in predictive pursuit to future target motion with unpredictable velocity.

Similar to predictive tracking for target motion with randomized timing (Barnes and Collins 2011; Collins and Barnes 2009), the history effects of previous trials on predictive pursuit was observed, and the effect of each previous trial declined progressively as the trial retracted into the past. These similar results suggest that predictive pursuit systems use a common strategy of weighted averaging of the information from previous trials to track the randomized velocity and timing of target motion. Of the retinal and extraretinal signals involved in smooth pursuit, the latter include the effects of volition, attention, and expectation, as well as memory of the target and eye velocity (Leigh and Zee 2015). Extraretinal signals can generate predictive pursuit that is inconsistent with visual feedback (retinal input) near the timing of target reversal (Boman and Hotson 1992). According to a model of smooth pursuit described by previous studies, a continual estimation of the timing is constructed by weighted averaging of stimulus history and retained in a form of working memory to initiate predictive pursuit (Barnes and Collins 2011; Fukushima et al. 2013). The memory of target velocity required for predictive pursuit can be held up to six variations (Barnes and Schmid 2002; Collins and Barnes 2005) and can be retrained for 14 s (Chakraborti et al. 2002). Given that it takes several trials for the predictive eye velocity to reach a saturation level (Barnes and Schmid 2002; Chakraborti et al. 2002; Collins and Barnes 2005), storing and continuously averaging information about the target velocity can be used to determine the scale of predictive pursuit to the randomized velocity.

Our findings showed that not only the stimulus history but also the behavioral history influences predictive pursuit to target motion with randomized velocity. For predictive pursuit to target motion with randomized timing, the influence of stimulus history has been demonstrated by multiple regression analyses (Barnes and Collins 2011, 2015; Collins and Barnes 2009) and a node method in which the sequential effect of the stimulus history is assesses by comparing the predictive pursuit responses sorted based on a characteristic of previous trials (Heinen et al. 2005; de Hemptinne et al. 2010, 2013; Kowler et al. 1984). Additionally, one of these studies has provided an evidence that predictive pursuit is influenced by the stimulus history but not the behavioral history (Collins and Barnes 2009). On the other hand, there is a study indicating that the onset of predictive pursuit is significantly influenced by the behavioral history but not the stimulus history (Badler and Heinen 2006). From these studies, the influence of behavioral history on predictive pursuit of randomly timed stimuli is ambiguous, but the following evidence may contribute to explaining the influence of behavioral history on predictive pursuit to the randomized velocity in this study. For example, although observers can generate predictive pursuit simply after observing a moving target (Barnes et al. 1997, 2000), it has been demonstrated that the velocity of predictive pursuit is greater after active pursuit than after passive observation (Burke and Barnes 2008b). Similar effects have been observed in eye motion to perturbations of target motion, the eye velocity induced by perturbations is greater when smooth pursuit was performed in previous trials than when it was not (Tabata et al. 2008). Taken together, execution of smooth pursuit affects the velocity of smooth pursuit in subsequent trials, which in turn may affect the control of predictive pursuit as the behavioral history. Intriguingly, while the fixed effects of the stimulus history of the n-2 and n-3 trials were significant, that of the most recent trial (n-1) were not. A similar result has been observed in predictive pursuit of randomly timed stimuli (Collins and Barnes 2009). In contrast, the fixed effects of the behavioral history appeared to correspond to the newness of the memory. Such a difference suggests that the information of sensory input (stimulus) and motor output (behavior) in the past time sequences have partially different influences on predictive pursuit.

Although the stimulus and behavioral history are not completely distinct, it may be useful to consider the differences in their influences on predictive pursuit from a neurophysiological perspective. The supplementary eye field (SEF) is known to be involved in predictive pursuit (de Hemptinne et al. 2008; Missal and Heinen 2004). Indeed, adding transcranial magnetic stimulation (TMS) to the SEF at the timing of target reversal increases eye velocity in the opposite direction, but adding TMS in the middle of the cycle does not affect eye velocity (Gagnon et al. 2006). Since the SEF is directly projected from the DLPFC (Wang et al. 2005), where the target information is likely to be stored in the form of working memory (Burke and Barnes 2008a; Schmid et al. 2001), it is considered to be essential for the execution of predictive pursuit based on the information obtained from previous trials. However, the SEF contains only a small portion of the neurons directly involved in generating motor responses (Fukushima et al. 2004; Shichinohe et al. 2009), indicating that the SEF plays a premotor role in predictive pursuit (de Hemptinne et al. 2008). The FEF, the major output center of smooth pursuit, receives signals from the SEF and generates predictive pursuit (Fukushima et al. 2013; Leigh and Zee 2015). In addition, the FEF modulates the smooth pursuit gain based on motor output in the past time sequences (Darlington et al. 2018). Altogether, the signal that generates predictive pursuit based on the working memory input via the SEF may be modulated by gain control based on behavioral history in the FEF.

In summary, our findings expand previous studies related to predictive pursuit. Predictive pursuit to target motion with randomized velocity showed history effects of previous trials, similar to predictive tracking for target motion with randomized timing (Barnes and Collins 2011, 2015; Collins and Barnes 2009). However, contrast to predictive pursuit of randomly timed stimuli, the influences of stimulus and behavioral history were found in this study. These results suggest that predictive pursuit systems can track the randomized velocity and timing of target motion using a common strategy of weighted averaging the information from previous trials, but the sources of information are partially distinct.

## Acknowledgements

This research was supported by JSPS KAKENHI Grant number 19K11460 and 18KK0286.

## References

Badler JB, Heinen SJ. Anticipatory movement timing using prediction and external cues. J Neurosci 26: 4519–4525, 2006.

Barnes GR. Cognitive processes involved in smooth pursuit eye movements. Brain Cogn 68: 309–326, 2008.

Barnes GR, Barnes DM, Chakraborti SR. Ocular pursuit responses to repeated, single-cycle sinusoids reveal behavior compatible with predictive pursuit. J Neurophysiol 84: 2340–2355, 2000.

Barnes GR, Collins CJS. The influence of cues and stimulus history on the non-linear frequency characteristics of the pursuit response to randomized target motion. Exp Brain Res 212: 225–240, 2011.

Barnes GR, Collins CJS. Influence of predictability on control of extra-retinal components of smooth pursuit during prolonged 2D tracking. Exp Brain Res 233: 885–897, 2015.

Barnes GR, Collins CJS, Arnold LR. Predicting the duration of ocular pursuit in humans. Exp Brain Res 160: 10–21, 2005.

Barnes GR, Grealy MA, Collins CJS. Volitional control of anticipatory ocular smooth pursuit after viewing, but not pursuing, a moving target: Evidence for a re-afferent velocity store. Exp Brain Res 116: 445–455, 1997.

Barnes GR, Schmid AM. Sequence learning in human ocular smooth pursuit. Exp Brain Res 144: 322–335, 2002.

Bland JM, Altman DG. Statistics notes: Calculating Correlation coefficients with repeated observations: Part 1— correlation within subjects. BMJ 310: 446, 1995.

Boman DK, Hotson JR. Predictive smooth pursuit eye movements near abrupt changes in motion direction. Vision Res 32: 675–689, 1992.

Burke MR, Barnes GR. Brain and behavior: A task-dependent eye movement study. Cereb Cortex 18: 126–135, 2008a.

Burke MR, Barnes GR. Anticipatory eye movements evoked after active following versus passive observation of a predictable motion stimulus. Brain Res 1245: 74–81, 2008b.

Carl JR, Gellman RS. Human smooth pursuit: Stimulus-dependent responses. J Neurophysiol 57: 1446–1463, 1987.

Chakraborti SR, Barnes GR, Collins CJS. Factors affecting the longevity of a short-term velocity store for predictive oculomotor tracking. Exp Brain Res 144: 152–158, 2002.

Collins CJS, Barnes GR. Scaling of smooth anticipatory eye velocity in response to sequences of discrete target movements in humans. Exp Brain Res 167: 404–413, 2005.

Collins CJS, Barnes GR. Predicting the unpredictable: Weighted averaging of past stimulus timing facilitates ocular pursuit of randomly timed stimuli. J Neurosci 29: 13302–13314, 2009.

Darlington TR, Beck JM, Lisberger SG. Neural implementation of Bayesian inference in a sensorimotor behavior. Nat Neurosci 21: 1442–1451, 2018.

Fiehler K, Brenner E, Spering M. Prediction in goal-directed action. J Vis 19, 2019.

Fukushima J, Akao T, Takeichi N, Kurkin S, Kaneko CRS, Fukushima K. Pursuit-related neurons in the supplementary eye fields: Discharge during pursuit and passive whole body rotation. J Neurophysiol 91: 2809–2825, 2004.

Fukushima K, Fukushima J, Warabi T, Barnes GR. Cognitive processes involved in smooth pursuit eye movements: Behavioral evidence, neural substrate and clinical correlation. Front Syst Neurosci 7: 1–28, 2013.

Gagnon D, Paus T, Grosbras MH, Pike GB, O’Driscoll GA. Transcranial magnetic stimulation of frontal oculomotor regions during smooth pursuit. J Neurosci 26: 458–466, 2006.

Goldman-Rakic PS. Chapter 16 Cellular and circuit basis of working memory in prefrontal cortex of nonhuman primates. Prog Brain Res 85: 325–336, 1991.

Heinen SJ, Badler JB, Ting W. Timing and velocity randomization similarly affect anticipatory pursuit. J Vis 5: 493–503, 2005.

de Hemptinne C, Barnes GR, Missal M. Influence of previous target motion on anticipatory pursuit deceleration. Exp Brain Res 207: 173–184, 2010.

de Hemptinne C, Ivanoiu A, Lefèvre P, Missal M. How does Parkinson’s disease and aging affect temporal expectation and the implicit timing of eye movements? Neuropsychologia 51: 340–348, 2013.

de Hemptinne C, Lefèvre P, Missal M. Neuronal bases of directional expectation and anticipatory pursuit. J Neurosci 28: 4298–4310, 2008.

Jacobs RJ. Visual resolution and contour interaction in the fovea and periphery. Vision Res 19: 1187–1195, 1979.

Jarrett CB, Barnes GR. Volitional scaling of anticipatory ocular pursuit velocity using precues. Cogn Brain Res 14: 383–388, 2002.

Kao GW, Morrow MJ. The relationship of anticipatory smooth eye movement to smooth pursuit initiation. Vision Res 34: 3027–3036, 1994.

Kowler E, Martins AJ, Pavel M. The effect of expectations on slow oculomotor control-IV. Anticipatory smooth eye movements depend on prior target motions. Vision Res 24: 197–210, 1984.

Kowler E, Rubinstein JF, Santos EM, Wang J. Predictive Smooth Pursuit Eye Movements. Annu Rev Vis Sci 5: 223–246, 2019.

Leigh RJ, Zee DS. The Neurology of Eye Movements. 5th ed. Oxford: Oxford University Press, 2015.

Matsuda K, Nagami T, Sugase Y, Takemura A, Kawano K. A widely applicable real-time mono/binocular eye tracking system using a high frame-rate digital camera. Lect Notes Comput Sci (including Subser Lect Notes Artif Intell Lect Notes Bioinformatics) 10271: 593–608, 2017.

Missal M, Heinen SJ. Supplementary Eye Fields Stimulation Facilitates Anticipatory Pursuit. J Neurophysiol 92: 1257–1262, 2004.

Miyamoto T, Miura K, Kizuka T, Ono S. Properties of smooth pursuit and visual motion reaction time to second-order motion stimuli. PLoS One 15: e0243430, 2020.

Ono S, Miura K, Kawamura T, Kizuka T. Asymmetric smooth pursuit eye movements and visual motion reaction time. Physiol Rep 7: 1–8, 2019.

Schmid A, Rees G, Frith C, Barnes G. An fMRI study of anticipation and learning of smooth pursuit eye movements in humans. Neuroreport 12: 1409–1414, 2001.

Shichinohe N, Akao T, Kurkin S, Fukushima J, Kaneko CRS, Fukushima K. Memory and Decision Making in the Frontal Cortex during Visual Motion Processing for Smooth Pursuit Eye Movements. Neuron 62: 717–732, 2009.

Tabata H, Miura K, Kawano K. Trial-by-trial updating of the gain in preparation for smooth pursuit eye movement based on past experience in humans. J Neurophysiol 99: 747–758, 2008.

Tanaka M, Lisberger SG. Regulation of the gain of visually guided smooth-pursuit eye movements by frontal cortex. Nature 409: 191–194, 2001.

Wang Y, Isoda M, Matsuzaka Y, Shima K, Tanji J. Prefrontal cortical cells projecting to the supplementary eye field and presupplementary motor area in the monkey. Neurosci Res 53: 1–7, 2005.

